# Low dose of a non-urea selective GIRK channel activator improves hippocampal-dependent synaptic plasticity and memory disrupted by amyloid-β oligomers

**DOI:** 10.1101/2024.11.05.622060

**Authors:** Jaime Mulero-Franco, Raquel Jimenez-Herrera, Ana Contreras, Souhail Djebari, Lydia Jiménez-Díaz, Juan D. Navarro-López

## Abstract

Increased neural activity characterizes early Alzheimer’s disease (AD), serving as a prognostic indicator for disease progression and cognitive decline. Mechanisms that drive this hyperactivity and their behavioral effects remain mostly unrevealed, although normalizing altered excitability levels has been shown to reverse cognitive impairment in early AD, both in animals and humans. Soluble amyloid-*β* oligomers (oA*β)* primary accumulate in limbic regions like hippocampus and induce neuronal hyperexcitability and subsequent cognitive deficits by impairing ion channel’s function. Indeed, G protein-gated inwardly rectifying K+ (GIRK) channels -that control neuronal excitability-are greatly affected and their selective pharmacological activation has already been shown very effective to counteract oA*β*-induced hyperexcitability and hippocampal dysfunction. However, GIRK gain-of-function in healthy animals disrupts learning, memory and underlying synaptic plasticity, greatly limiting its therapeutic potential in preclinical asymptomatic early AD patients. Therefore, GIRK-based pharmacological treatment needs further investigation to overcome these limitations. Here we tested two doses of a novel, more potent, and neuronal selective GIRK activator, VU0810464, in healthy and early oA*β*_1-42_-generated AD male and female mice. Both doses normalized hippocampal synaptic plasticity (long-term potentiation, LTP) and associated spatial object location memory (OLM) without sex dimorphism in AD animals. However, in healthy mice, low VU0810464 dose did not significantly alter LTP and OLM, whereas the high dose disrupted both. Our results suggest that the precise tuning of neural excitability with low dosing of VU0810464 might be a promising strategy to safely treat and prevent hippocampal overexcitation and upstreaming memory deficits in early preclinical asymptomatic phases of AD.

## 1. Introduction

Alzheimer’s disease (AD), a condition that accounts for 60-80% of the 55 million people with dementia [1], increases its prevalence significantly with age, doubling every 5 years from the ages of 50 to 80 [2]. AD exhibits notable sex differences as almost two-thirds of patients with AD are female (Mielke et al., 2014). The progression of AD likely spans a period of 20–30 years, so therapeutic interventions to prevent or delay symptoms should be focused on very early stages, even asymptomatic, when these sex differences are not evidenced yet [3].

AD is characterized by accumulation of amyloid-*β* (A*β*) plaques and neurofibrillary tangles (NFTs) consisting of hyperphosphorylated tau protein in the brain. However, one of the primary events in AD is thought to be the aggregation of A*β* in limbic regions like hippocampus [4], promoting the migration and deposition of tau in the majority of older individuals, contributing to the onset of AD expression [5–7]. In early stages of AD, the oligomeric soluble forms of A*β* (oA*β*) are one of the most predominant pathogenetic factors [8]. The presence of oA*β* is linked to impaired inhibitory GABAergic interneuron function [9], abnormal glutamate release and reuptake, and dysfunction of ion channels [10, 11]. Its neurotoxic effects induce hippocampal dysfunction due to neuronal hyperexcitability, leading to excitatory/inhibitory (E/I) neurotransmission imbalance, synaptic plasticity disturbances, disrupted oscillatory activity, and cognitive deficits [12]. Therefore, neuronal hyperexcitability induced by oA*β* emerges as an interesting therapeutic target to be explored in early AD, as normalizing excitability levels has been shown to be beneficial not only in transgenic [13] and non-transgenic [14] models of AD, but also in human patients [15].

In the hippocampus, G protein-gated inwardly rectifying potassium (GIRK) channels are tetramers composed by GIRK1/2 subunits which mediate the inhibitory G protein (G_i/o_)-coupled effects of neurotransmitters including GABA, adenosine, or 5-HT among others, as the opening of the channels lead to a K^+^ efflux and neuron hyperpolarization [16]. GIRK activity impairment has been linked to multiple disorders [16, 17], including AD [18]. In the dorsal hippocampus, they are constitutively active, and their bidirectional regulation by *gain-* or *loss-of-function* lead to cognitive impairments in mice [19–22] through neuronal excitability [18, 23], synaptic [19, 21, 24] and network [25, 26] dysfunctions and disturbance of the E/I balance. These data point to GIRK channels as a promising therapeutic target for restoring E/I equilibrium [16, 27], although with a narrow regulation range to avoid deleterious effects on hippocampal performance [19]. Hippocampal GIRK channels were associated to AD amyloid pathology more than one decade ago by our group [18]. Since then, growing evidence has shown them as potential target to counteract A*β*-induced hyperexcitability and associated impairments at different levels of complexity, from molecular to synaptic and behavioral [14, 28–35], and additional mechanisms involving GIRK channels in A*β*-induced dysregulations through a signaling cascade mediated by PrPC and mGluR5 have been proposed [35]. Their activation with the selective GIRK1-containing opener ML297 in the hippocampus counteracts synaptic, network and behavioral deficits induced by oA*β* [14, 28, 29], although such drug has shown different pharmacodynamic and pharmacokinetic limitations, in addition to disruption of synaptic plasticity and subsequent learning and memory processes in healthy animals [19], both greatly limiting its potential use to prevent oA*β* effects in preclinical asymptomatic early AD patients.

Here, we use non-transgenic male and female mice of an early AD model generated by a single intracerebroventricular (*icv*.) injection of oA*β* to test the effects of high and low doses of a novel GIRK activator with enhanced potency and selectivity for neuronal GIRK1/2 channels, VU0810464 [36], on hippocampal long-term synaptic plasticity (LTP) and dorsal hippocampal-dependent memory processes. Our data show that both VU0810464 concentrations do attenuate cognitive impairments found in the early AD model by restoring synaptic plasticity. However, in healthy control animals, high VU0810464 concentration significantly blocked synaptic plasticity and impaired hippocampal-dependent memory, as previous findings with the GIRK channel opener ML297, while the low dose did not induce any of these alterations. Hence, our results suggest this latter low dose as a potential treatment in models of early AD, and may also offer potential preventive benefits in the preclinical stages of this dementia [37].

## 2. Materials and methods

### 2.1. Animals

70 C57BL/6 adult mice, comprised of 33 females and 37 males (3 to 6 months of age; 20 to 35 g; bred in-house) were housed in the Animal House Facilities of the Universidad de Castilla-La Mancha (Ciudad Real, Spain). This age range falls within the scope of mature adulthood [38]. Animals were housed in groups of 4 to 5 per cage before surgery, with environmental conditions strictly controlled to maintain a temperature of 21 ± 1°C and a humidity level of 50 ± 7%. They had access to food and water *ad libitum* and environmental enrichment.

All experimental procedures were reviewed and approved by the Ethical Committee for Use of Laboratory Animals of the University of Castilla-La Mancha (PR-2021-12-21 and PR-2023-25) and conducted in accordance with European Union guidelines (2010/63/EU) and spanish regulations governing the use of laboratory animals in chronic experiments (RD 53/2013 on the care of experimental animals: BOE 08/02/2013).

### 2.2. Surgery for intracerebroventricular drug injection

Drugs used in this study were administered through *icv.* injections, that mainly target the dorsal hippocampus as already extensively validated by our group and others [28, 39, 40]. For *icv*. administration, animals were surgically implanted with a blunted, stainless steel, 26-G guide cannula (Plastics One, US) directed towards the lateral ventricle (0.5 mm posterior to bregma, 1.0 mm lateral to midline, and 1.8 mm below the brain surface) (Fig. 1A).

**Figure 1.**
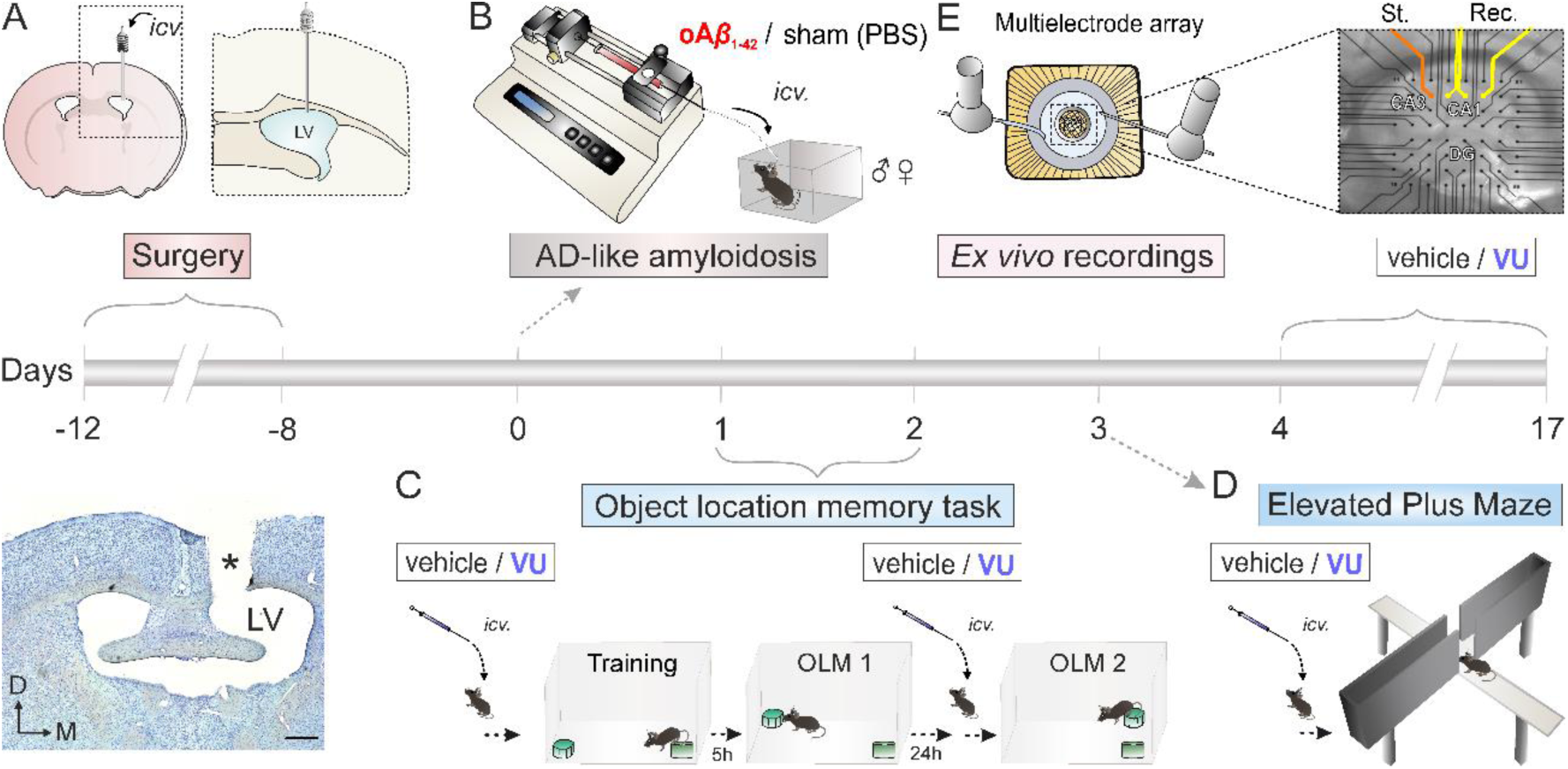
Experimental design showing the timeline for recordings and behavioral testing. **(A)** A guide cannula was implanted for *icv.* drug administration on the left ventricle, which was later histologically verified (asterisk, *, indicates cannula’s location). Scale bar: 500 μm. (**B)** On day 0, after surgery recovery, a single *icv.* injection of oA*β*_1–42_ was administered to set up a non-transgenic model of early AD-like amyloidosis in the dorsal hippocampus. Control sham animals were *icv*. injected with PBS. **(C-D)** Selective GIRK channel activator VU0810464 (VU) or its vehicle were administered daily 30 min before behavioral testing. **(C)** On day 1 post-oA*β* or sham PBS *icv.* delivery, animals received an *icv*. injection of low or high dose of VU or its vehicle before the training session for the Object Location memory (OLM) task. Five hours after training, a short-term memory test session (OLM 1) was performed. On day 2 post amyloidosis induction by oA*β*, animals received another *icv*. injection of low or high dose of VU before a test session for long-term memory retrieval (OLM 2) was conducted. **(D)** On day 3 post-oA*β* injection, animals received the corresponding VU treatment and anxiety levels were measured using the Elevated Plus Maze. **(E)** *Ex vivo* hippocampal LTP was studied using multi-electrode arrays (MEAs) electrophysiology in hippocampal horizontal slices (days 4–17 post-oA*β* injection) adding VU at low or high doses to the perfusion system. Representative location of stimulation (St., orange) and recording (Rec., yellow) electrodes in a hippocampal horizontal slice. *icv*., intracerebroventricular; LV, lateral ventricle; D, dorsal, M, medial.

Briefly, mice were anesthetized with 4% isoflurane (#13400264, ISOFLO; Proyma S.L., Spain) delivered using a calibrated R580S vaporizer (RWD Life Science, China; flow rate: 0.5 L/min O_2_). Following induction, a constant delivery of 1.5-2% isoflurane was used to maintained anesthesia. Buprenorphine was administered intramuscularly as analgesic during and after surgery (0.01 mg/kg; #062009, buprenodale; Albet, Spain) and a healing cream (#732438, Blastoestimulina®; Almirall, Spain) was applied to promote recovery and minimize suffering and discomfort experienced by the animals. Animals were allowed a recovery period of at least one week to ensure their well-being before starting any experimental procedures.

Injections were conducted in alert freely moving mice (Fig. 1B) using a motorized Hamilton syringe at a rate of 0.5 μl/min through a 33-G injection cannula, which was 0.5 mm longer than the chronically implanted guide cannula and inserted inside it, following established protocols [40]. Specific temporal parameters for drug *icv*. injections are described for each experimental condition (see Fig. 1).

Animals prepared as described above were used for behavioral testing and *ex vivo* electrophysiological recordings (Fig. 1). As a preliminary step, to evaluate the potential side effects of *in vivo* administration of two different low dosages of the selective GIRK activator VU0810464, and therefore choose the most appropriate one for our present study, a pilot experiment was conducted (Fig. S1, Supplemental Information S1). Behavioral assessments in this pilot included Barnes maze testing, Object Location Memory (OLM) task, Elevated Plus Maze (EPM), stereotyped behaviors test, and Rotarod to determine the effects of two different VU0810464 concentrations on spatial memory and locomotion in healthy animals (Fig. S1).

Following dose selection, the main study was conducted in a non-transgenic early AD-like amyloidosis mouse model using the selected VU0810464 concentration from the pilot study, as well as a higher concentration similar to the one previously used for *in vivo* experiments with the selective GIRK activator ML297 [19]. Behavioral assessments included OLM and EPM testing (Fig. 1C and 1D, respectively). *Ex vivo* field excitatory postsynaptic potential (fEPSP) recordings were also performed to evaluate the efficacy of VU0810464 in the early AD-like mice and to further confirm the safety of the chosen doses in control healthy animals (Fig. 1E).

### 2.3. Non-transgenic early AD mouse model

This model has been extensively characterized and validated by our group and others to study early AD-like neuropathology at molecular, synaptic, network and behavioral levels by resembling early acute amyloidosis in dorsal hippocampal regions [14, 28, 29, 40, 41]. In order to generate the non-transgenic early AD mouse model, once fully recovered from surgery procedures, awake freely moving animals underwent a single 3 μL *icv*. injection of oA*β*_1-42_ (1 μg*/*μL; 0.15 μg/g; monomeric A*β*_1-42_ #ab120301; Abcam, UK; Fig. 1B) as previously described [40]. To obtain soluble oligomers of the 1-42 fragment of the amyloid peptide (oA*β*_1-42_), monomeric soluble A*β*_1-42_ was dissolved in PBS and incubated 6 hours at 37°C [42, 43]. Concentration of oA*β*_1-42_ was selected in accordance with previous studies [40] and presence of oA*β*_1-42_ in hippocampal tissue was also checked by western blot after mice decapitation and fresh hippocampi extraction as previously shown [40]. Control sham animals (referred to as sham healthy mice) were *icv*. injected with 3 μL of PBS.

### 2.4. Pharmacological modulation of GIRK channels activity. Drug dosing

To pharmacologically modulate GIRK channels activity, we employed VU0810464 (#HY-127106; MedChemExpress, US), a selective non-urea-based activator of GIRK1/2 channels (Fig. 1C-E). This compound has demonstrated superior blood-brain barrier penetration and selectivity for neuronal GIRK channels compared to previously reported GIRK channel activators such as ML297 [44]. Also, VU0810464 displays increased potency as a GIRK1/2 activator and improved brain distribution in *in vivo* pharmacokinetic studies in mice [44]. For *ex vivo* experiments, two concentrations of VU0810464 dissolved in DMSO (0.2-0.4% DMSO in bath conditions) underwent through evaluation, a high concentration (10 µM) based on previous studies [44] and a lower one: 5 µM (Fig. 1E). Similarly, for *in vivo* experiments, two concentrations of VU0810464 (high:1.5 mM (0.07µg/g) and low: 0.75 mM (0.035µg/g)) were selected for *icv.* administration. The low and high doses were chosen based on results of the exhaustive pilot experiments showed in supplementary material and supplementary Fig. S1 and previous experiments using other GIRK opener [19]. For *in vivo* administration, VU0810464 was dissolved in PBS with the help of 10% *β-*hydroxypropyl cyclodextrin (HPβCD) as vehicle based on previous reports [44].

In summary, the same six experimental groups were analyzed both for *ex vivo* and *in vivo* experiments, i.e., three treatments (high dose of VU0810464, low dose of VU0810464, and vehicle) were tested in two conditions: healthy (control sham mice *icv*.-injected with PBS) and pathological (AD-like animals oA*β*_1-42_-injected). In order to clarify descriptions and figures, in all panels, VU0810464 and oA*β*_1-42_ have been abbreviated as VU and oA*β* respectively. All experiments were conducted using a blinded design, so that the researcher did not know the treatment of each animal to avoid possible bias.

### 2.5. Object location memory

OLM test was employed to assess hippocampal-dependent short-and long-term spatial memory [40, 45] in our AD mouse model (Fig. 1C). Our experimental design allowed us to evaluate the effect of VU0810464 on short-and long-term spatial memory retrieval.

As shown in Fig. 1B, on Day 0, mice were injected with either oA*β*_1-42_ (AD-like mice) or PBS (control sham healthy animals) and, 1 h later, they were habituated to a 49 × 49 cm square box for a duration of 10 min, with visual cues positioned around the room to offer spatial context. The following day (Day 1; Fig. 1C), AD and sham healthy mice received a single *icv*. injection of either VU0810464 or its vehicle, and 30 min later a training session was performed. In this session, two objects were placed within the chamber, and mice were allowed to freely explore them for 10 min. Following the training session, 5 h later, short-term memory retrieval assessment (OLM 1) was performed. During this test, one of the objects was moved to a novel location within the box. 24 h later, to evaluate long-term memory retrieval (Day 2; Fig. 1C), mice were *icv.* injected again with the corresponding treatment (VU0810464 or vehicle), and 30 min post-*icv*. injection, another memory test was conducted (OLM 2) to evaluate long-term memory, moving the same object to a third location. To eliminate any residual odor cues, both box and objects were cleaned with 70% ethanol between each mouse trial.

The time spent exploring each object was analyzed, and discrimination index (DI) was computed for training, OLM 1 and OLM 2 with the following formula: (Time exploring moved object – time exploring unmoved object)/Total exploration time. Exploration of the objects was defined as the amount of time mice were oriented toward an object with its nose within 1 cm of it. Animals with a total exploration time lower than 5 s were excluded from the analysis [45]. Data from the training session were used to ensure that animals did not show an initial preference for either side of the box, so animals with a DI ≤ 0.2 were excluded from the rest of the analysis.

### 2.6. Elevated plus maze

After OLM assessment, the Elevated Plus Maze (EPM) was used to evaluate anxiety-like behaviors (Day 3; Fig. 1D). EPM comprises a methacrylate platform in a cross shape, featuring two open arms (65 × 6 cm) without walls and two enclosed arms (65 × 6 cm) with 15-cm-high opaque walls. These arms are arranged at right angles to each other, centered around a platform (6.3 × 6.3 cm), and elevated 40 cm above the floor. Mice exposed to different concentrations of VU0810464 were introduced in the center of the maze and allowed to freely explore the platform during a single 5-min session. Number of entries in open arms normalized by the total number of entries in both arms was analyzed as an indicator of anxiety-like behavior. Additionally, total distance covered was evaluated as a measurement of locomotor activity. Sessions were recorded and analyzed with the Smart video tracking software (Panlab, Spain).

### 2.7. Ex vivo fEPSP recordings

Electrophysiology recordings were conducted in horizontal hippocampal slices obtained from AD-like or sham mice following previously described procedures [40]. Therefore, hippocampal slices were obtained 4 to 17 days after *icv.* injection of oA*β*_1_-_42_ or PBS (sham controls), a timeframe during which the effects of a single oA*β*_1-42_ *icv.* injection have been shown to persist [40] (Fig. 1B, E). Mice did not receive *icv*. VU0810464 before slicing, as the drug was perfused in the slices. Briefly, mice were anesthetized with halothane (Fluothane; AstraZeneca, UK) and subsequently decapitated. The brain was immediately taken out and quickly immersed in an oxygenated (95% O2 – 5% CO2) ice-cold “cutting” solution containing (in mmol/L; all from Sigma, US): 40 NaCl (#S9888), 10 glucose (#G8270), 150 sucrose (#84100), 1.25 NaH2PO4 (#S8282), 26 NaHCO3 (#S6014), 4 KCl (#P3911), 0.5 CaCl2 (#499609), and 7 MgCl2 (#208337). The brain was excised and affixed onto the stage of the vibratome (7000smz-2; Campden Instruments, UK) allowing the blade to cut through both hemispheres at an angle of 15-20° from their horizontal planes. Slices were cut to a thickness of 300 µm containing the dorsal hippocampi. They were then incubated for 1.5 h to 2 h at room temperature (22-23°C) in an oxygenated artificial cerebrospinal fluid (aCSF) containing (in mmol/L; all from Sigma, US): 125 NaCl (#S9888), 3.5 KCl (#P3911), 2.4 CaCl2 (#499609), 1.3 MgCl2 (#208337), 26 NaHCO3 (#S6014), 25 glucose (#G8270), and 1.2 NaH2PO4 (#S8282).

For electrophysiological *ex vivo* fEPSP recordings, two set-ups, each compromising a multi-electrode array (MEA2100-Mini-System) pre-amplifier and a filter amplifier (gain 1100× or 550×) were run simultaneously. Both using a data acquisition card governed by MC_Experimenter V2.20.0 software. Every single slice was moved to each MEA recording chamber (MEA60; Multi Channel Systems, Reutlingen, Germany), which was continuously perfused with aCSF (flow rate 2 mL/min) and kept at 32°C. MEA chambers, that were placed on the platform of an inverted MEA-VMTC-1 video microscope, consisted of 60 extracellular electrodes, with a distance of 200 μm between each electrode. Each individual electrode within the array had the capability to function either as a recording or a stimulation electrode. A nylon mesh was placed over the slice to guarantee optimal electrical contact between the slice surface and the electrode array. Stimulation was achieved using a stimulus generator unit incorporated into the headstage (Multi Channel Systems, Germany), which delivered biphasic current pulses to a designated electrode situated within the Schaffer collateral pathway of the hippocampus.

fEPSPs were recorded as previously described [40]. Briefly, the *stratum radiatum* of the CA1 subfield was recorded by all the remaining electrodes of the array simultaneously (Fig. 1E). After an equilibration period of at least 20 min inside the MEA chamber, responses of the Schaffer collateral pathway were recorded. Each slice underwent a 15-min period of stabilization followed by 30 min of treatment with either VU0810464 or its vehicle before LTP induction. The drug was then perfused for an additional 5 min post-HFS before being washed out.

For LTP induction, a high-frequency stimulation (HFS) protocol was used, consisting of three 1-second 100-Hz trains delivered at 10-min intervals between trains. fEPSP amplitude measurements taken 15 min before HFS, in the presence of the treatment, were used as the baseline (BL), while measurements 60 min after HFS were used to assess LTP induction.

Data were processed by using the Multichannel Analyzer software (Version 2. 20.0). As synaptic responses lacked population spikes, the amplitude (peak-to-peak value in mV during the rise-time period) was quantified for successively evoked fEPSPs. All values are represented as mean ± SEM with *n* indicating number of slices. Within each slice, data from 3 different recording electrodes were used.

### 2.8. Statistical analysis

Data were expressed as the mean ± SEM. Three-or two-way ANOVA were performed, using treatment, sex and time as variables when appropriate, and followed by Tukey’s *post-hoc* examination to compare differences between subgroups. Statistical significance was established at *p* < 0.05. All statistical analyses were conducted using SPSS software v.24 (RRID:SCR_002865; IBM, US) and GraphPad Prism software v.8.3.1 (RRID:SCR_002798; Dotmatics, US) and detailed in the text or in Table 1. Final figures were made using CorelDraw X8 Software (RRID:SCR_014235; Corel Corporation, Canada).

**Table 1.**
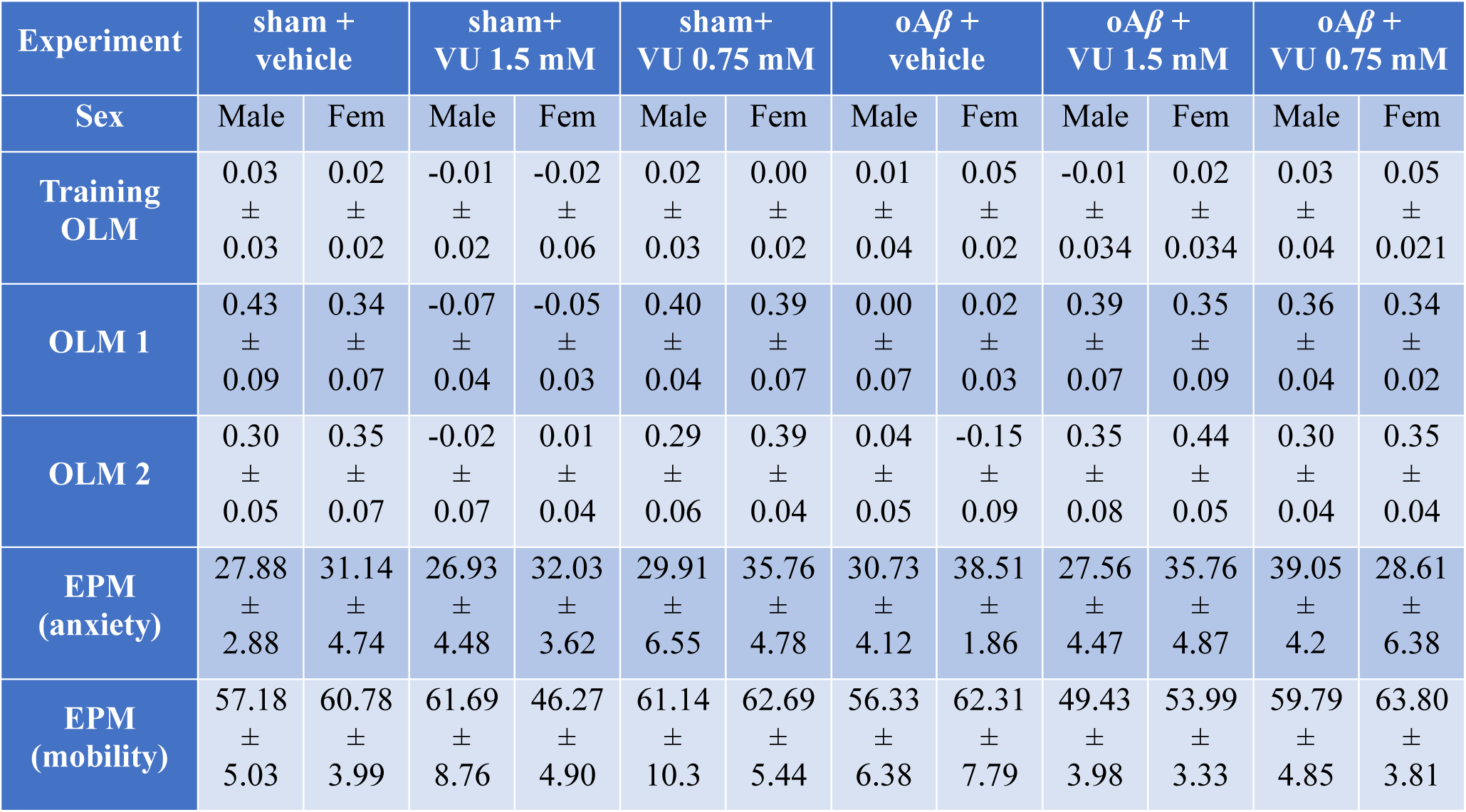
Statistical results for each experimental group on each behavioral task. . Specific Discriminations Index (DI) data for Object Location Memory (OLM) test during training, OLM 1 and OLM 2, as well as percentage of entries in the open arms test and total distance traveled in the Elevated Plus Maze (EPM). All data are represented separately for male and female.

## 3. Results

### 3.1. Low VU0810464 concentration improved LTP in oA*β*-treated mice without effects on healthy control animals

It has been previously shown that GIRK *gain-of-function* using ML297, a selective GIRK1-containing activator, induces deleterious effects on hippocampal CA3-CA1 synaptic plasticity and associated mechanisms of learning and memory [19]. However, in a model of amyloidosis, ML297 improves synaptic plasticity and memory [14, 28]. Here, we wondered whether VU0810464, a compound with presumable improved potency and selectivity compared to ML297 [44], would have similar effects in an early AD model but, which is more important, without adverse effects observed in healthy animals [19]. To this end, *ex vivo* electrophysiological recordings in a multielectrode array were obtained from hippocampal slices of animals 4 to 17 days post-*icv*. injection with oA*β*_1-42_ or PBS (Fig. 2A).

**Figure 2.**
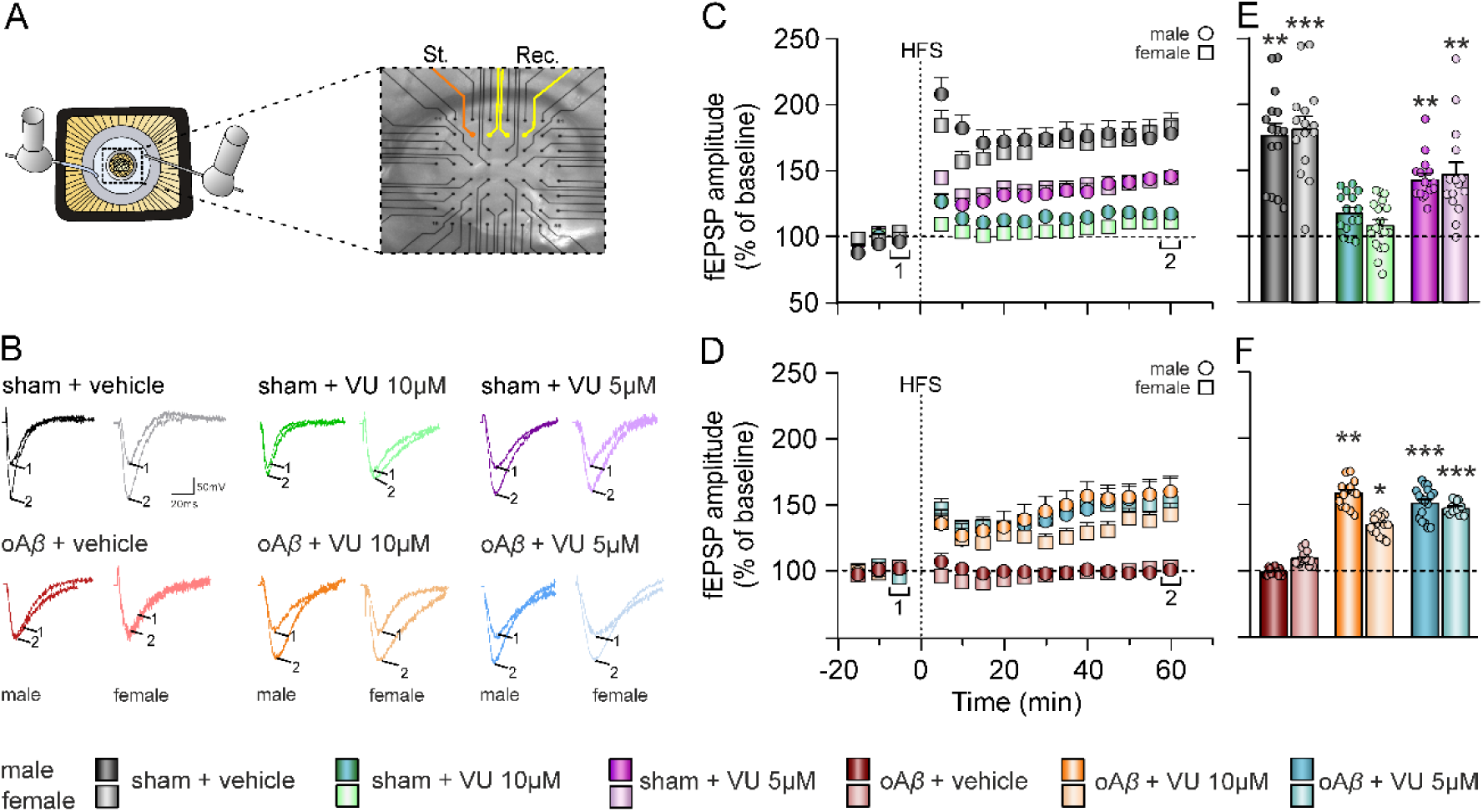
GIRK channels activator VU0810464 rescues long-term potentiation in oA*β*-treated mice. **(A)** Multi-electrode arrays (MEAs) electrophysiology was performed to study the effect of low or high concentration of VU0810464 (VU) on hippocampal LTP after oA*β*_1–42_ injection. VU or its vehicle were perfused in the recording chamber. **(B)** Representative averaged (n = 5-6) traces of fEPSPs recorded in the CA1 area, collected during the baseline (1) and ≈60 min post-HFS (2). **(C, D)** Time course of the LTP evoked in the CA1 area after a HFS protocol in hippocampal slices from **(C)** sham + vehicle, sham + VU 10µM and sham + VU 5µM and from **(D)** oA*β* + vehicle, oA*β* + VU 10µM or oA*β* + VU 5µM treated mice. **(E, F)** Bars illustrate mean ± SEM fEPSPs amplitude of the last 10 min of the recording from the same groups illustrated in C and D, respectively. * *p* < 0.05, ** *p* < 0.01, *** *p* < 0.001 *vs*. sham + VU 10µM (D) or *vs*. oA*β* + vehicle (F). St., stimulation electrode; Rec., recording electrode.

To test VU0810464 in healthy control animals, we began by assessing the effects on hippocampal CA1 synaptic plasticity of VU0810464 perfused at two different concentrations, high (10 µM) and low (5 µM), or its vehicle during 30 min pre-HFS and 5 min post-HFS (Fig. 2). In slices from sham mice perfused with vehicle, HFS protocol applied at Schaffer collaterals resulted in a 178.2 ± 11.5 % (*p* = 0.0002; male, black circles) and 184.5 ± 12.6 % (*p* < 0.0001; female, grey squares) potentiation of the response evoked in CA1 that lasted at least 1 hour, indicating successful LTP induction and maintenance (Fig. 2B, C, E). Notably, only the low VU0810464 dose evoked a successful significant HFS-dependent LTP (145.7 ± 6.5 %, *p* = 0.0004, in male and 147.2 ± 11.4 %, *p* < 0.0001, in female; pink circles and squares respectively), while the high dose hindered this plasticity process to pre-HFS values (118.8 ± 4.4 %, *p* = 0.1097, in male and 108.1 ± 6.8 %, *p* = 0.7459, in female; green circles and squares respectively; Fig. 2B, C, E), indicating that a lower concentration of VU0810464 does not impair synaptic plasticity in sham healthy subjects.

On the other hand, in slices obtained from AD-like oA*β*-injected animals perfused with vehicle, LTP was found blocked, and post-HFS response was approximately the same as baseline levels (97.0 ± 9.3 %, *p* > 0.9999, in male and 103.3 ± 5.3 %, *p* > 0.9999, in female; red circles and squares). This inhibition of synaptic plasticity by oA*β* was observed immediately after the HFS and persisted for at least 60 min afterwards (Fig. 2B, D, F), suggesting an alteration of the processes of LTP induction as described previously by our group [28, 40].

Next, we wondered whether activation of GIRK channels with VU0810464 in oA*β* early AD mice could normalize excitability levels and improve synaptic plasticity, both processes previously shown increased and hindered respectively by oA*β*_1-42_ [14, 28]. Slices were perfused with low or high concentrations of VU0810464 and, in all cases, LTP could be generated and maintained for more than 1 hour (oA*β*_1-42_ + VU0810464 10 µM (high dose, orange symbols): 160.1 ± 11.8 %, *p* = 0.0086, in male and 142.3 ± 9.1 %, *p* = 0.0128, in female; oA*β*_1-42_ + VU0810464 5 µM (low dose; blue symbols): 163.2 ± 9.1 %, *p* = 0.0008, in male and 152.8 ± 5.9 %, *p* < 0.0001, in female; Fig. 2B, D, F), suggesting that selective GIRK activation with low and high doses of the drug can restore successful induction of this type of synaptic plasticity previously hindered in AD-like oA*β*-injected animals.

Furthermore, when applying a three-way ANOVA to assess the effect of treatment taking into account the possible sex dimorphism, data showed a significant treatment effect due to VU0810464 administration (Fig. 2B-F; F_(5,174)_ = 33.03, *p* < 0.001) regardless of the sex (F_(1,174)_ = 1.026, *p* = 0.312). *Post-hoc* analysis revealed differences between control sham + vehicle and sham treated with VU0810464 10 µM (high dose) in both male (*p* = 0.0005) and female (*p* = 0.0001) mice indicating that the high dose of VU0810464 hindered LTP (as GIRK activator ML297 had previously shown [46]) in healthy animals regardless of sex. Also, *post-hoc* analysis between both concentrations of VU0810464 showed differences in male (*p* = 0.0056) and female (*p* = 0.0030) healthy mice indicating that reducing the VU0810464 concentration allows LTP induction and maintenance in control sham mice. Regarding oA*β*_1-42_ treated animals, *post-hoc* analysis revealed differences between oA*β* + vehicle and oA*β* treated with both concentrations of VU0810464 in male (10 µM, *p* = 0.0038; 5 µM, *p =* 0.0001) and female (10 µM, *p* = 0.01; 5 µM, *p* = 0.0001) mice.

Altogether, these results indicate that VU0810464 prevents LTP disruption in early AD-like A*β* amyloidosis mice without noticeable effects on healthy animals’ hippocampal plasticity and presumably associated memory, supporting the hypothesis that lowering dose could be a potential candidate for *in vivo* treatment in this pathology.

### 3.2. Low dose of VU0810464 counteracts spatial memory deficits caused by oA*β* without noticeable effects on healthy animals’ memory

Given that LTP has been shown to typically sustain hippocampus-dependent spatial learning [47], the OLM test was used to evaluate spatial short-and long-term hippocampal-dependent memory. Concentration for *in vivo* experiments were selected after an extensive and meticulous pilot study detailed in the supplementary material. We found the lowest concentration used in this pilot study (0.25 mM) to cause noticeable side effects on animals’ mobility, motor coordination, and stereotypic behaviors (Fig. S1). This suggests a possible phenomenon of hormesis, where very low and high doses of VU0810464 would induce undesirable effects, while beneficial doses lie in between [48]. This interesting phenomenon deserves further investigation in the future. However, *icv.* administration of 0.75 mM VU0810464 did not produce any of those deleterious effects in healthy animals (Fig. S1). So, 0.75 mM was chosen as low concentration of VU0810464 for *in vivo* memory testing, along with a higher concentration, 1.5 mM, similar to the one previously used for *in vivo* experiments with the selective GIRK activator ML297 [19], to further study which could successfully reverse the effects of oA*β* in our non-transgenic mice model of AD.

For OLM experiments, animals were *icv.* administered daily with VU0810464 or its corresponding vehicle throughout the duration of the experiments (i.e. days 1 and 2 post-*icv.* injection of oA*β*_1-42_ or sham PBS (Fig. 3A), as it has been shown that the drug is rapidly cleared *in vivo*, indicating that it is efficiently metabolized and eliminated [44]. Data from the OLM training session (Fig. 3A-C) showed that all animals spent equal amount of time exploring both objects (DI ≈ 0) which accounts for no preference for a specific object that could have influenced the later results. During both short-or long-term memory OLM tests (OLM 1 and OLM 2), three-way ANOVA showed no differences in the DI due to either sex within each treatment group (F_(1,57)_ = 0.002285, *p* = 0.962) or time (OLM 1 *vs.* OLM 2; F_(1,57)_ = 0.6066, *p* = 0.4393), indicating that males and females responded equally to the different treatments, and that the effect was consistent on both short-and long-term memory (Table 1).

**Figure 3.**
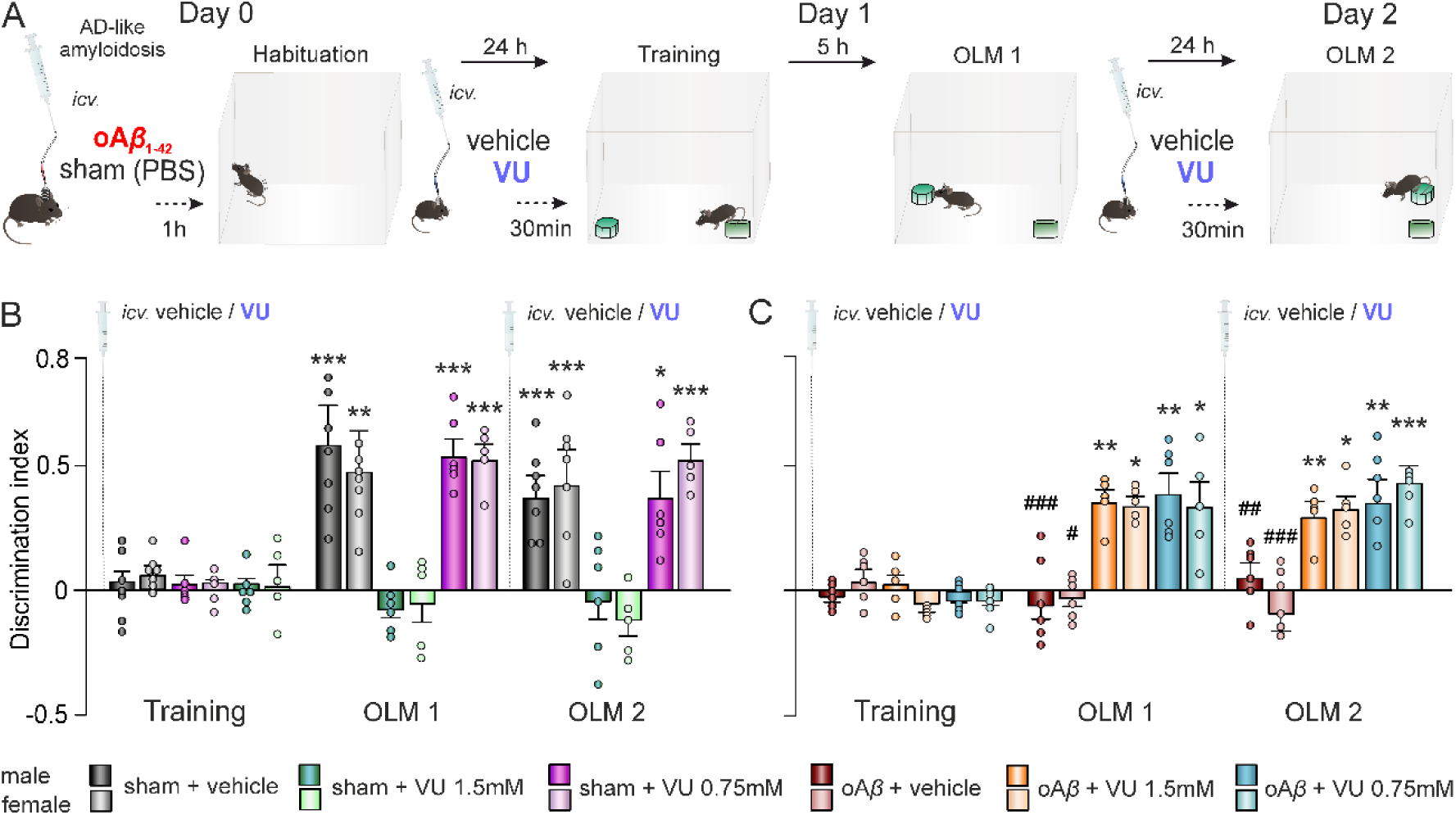
A low dose of VU0810464 improves spatial memory impaired by oA*β* without affecting healthy individuals. **(A)** An OLM test was performed, changing the location of one object between the training and each memory test (OLM 1 and OLM 2). oA*β*_1–42_ or sham (PBS) were administered *icv.* the day before training (day 0), and either vehicle or low or high doses of VU0810464 (VU) were *icv*.-injected daily 30 min before the behavioral task (days 1 and 2). **(B-C)** Discrimination index during training, OLM 1, and OLM 2 sessions for **(B)** sham + vehicle, sham + VU 0.75mM and sham + VU 1.5mM mice and **(C)** oA*β* + vehicle, oA*β* + VU 0.75mM and oA*β* + VU 1.5mM groups. Data are expressed as the mean ± SEM. * *p* < 0.05, ** *p* < 0.01, *** *p* < 0.001 *vs*. sham + VU 1.5mM (B) or *vs.* oA*β* + vehicle (C); # *p* < 0.05, ## *p* < 0.01, ### *p* < 0.001 comparing oA*β* + vehicle vs. sham + vehicle.

Interestingly, regarding the effect on sham healthy mice, the high dose of VU0810464 administered to sham animals (male, n = 6; female, n = 5) disrupted spatial memory (Fig. 3B) exhibiting a lower DI compared to vehicle in both tests. However, no significant effects were observed in either OLM tests with the low dose (male, n = 6; female, n = 5), as shown in Fig. 3B (Table 1). These results indicate that a low dose of VU0810464 does not disrupt spatial memory in sham healthy mice while the high dose does, in accordance with our previous *ex vivo* synaptic plasticity results.

On the other hand, when the effect of the GIRK activator VU0810464 was tested in early AD-like amyloidosis mice, a significant treatment effect of oA*β*_1-42_ was found (F_(5,57)_ = 34.93, *p* < 0.001). *Post-hoc* analysis revealed, as already reported by our group previously [40], that oA*β*_1-42_ + vehicle treated mice (male, n = 6; female, n = 6) exhibited a decreased DI relative to sham PBS-treated animals (male, n = 7; female, n = 7) (Fig. 3B, C) indicating impaired spatial memory induced by oA*β* in AD mice. Furthermore, both high (male, n = 5; female, n = 5) and low (male, n = 6; female, n = 5) doses of VU0810464 were able to counteract the short-(OLM 1 test) and long-term (OLM 2 test) memory deficits caused by oA*β*_1-42_ (Fig. 3C; Table 1) in accordance with our *ex vivo* synaptic plasticity results.

Overall, these results indicate that while both doses of VU0810464 were able to improve the memory deficits induced by early amyloidosis, the lower dose (0.75 mM) appeared to be a more efficient and safer concentration to prevent oA*β*_1-42_ deleterious effects as it does not affect sham healthy subjects. This might imply that a low dosage of the drug could be safely and preventively administered for AD models in prodromic asymptomatic stages.

### 3.3. VU0810464 did not induce anxiety-like behavior or locomotion impairments

To confirm that memory impairments found in AD mice were due to specific hippocampal alterations triggered by oA*β*_1–42_ injection, anxiety-like behavior and locomotion were assessed after OLM experiments. For that purpose, animals were challenged to the EPM task, a widely recognized ethologically-based test for anxiety in rodents, leveraging their innate aversion to open spaces and heights [49]. In addition, we could evaluate the possible GIRK-dependent decrease in anxiety-related behavior previously described [50].

Due to previous evidence of rapid VU0810464 clearance [44], oA*β*_1-42_ and sham *icv.-*injected mice received an additional *icv.* injection of the corresponding concentration of VU0810464 one day after OLM 2 testing (i.e., day 3 post-*icv.* injection of oA*β* or sham PBS) (Fig. 1D, 4A). Then, EPM was conducted 30 min after VU0810464 injection to evaluate this behavior.

Regarding stress-related behaviors, all animals displayed similar percentage of entries into open arms (Fig. 4B; treatment effect: F_(5,57)_ = 0.466, *p* = 0.80) regardless of sex (sex effect: F_(1,57)_ = 1.352, *p* = 0.2498), showing equivalent levels of anxiety (Table 1). Furthermore, distance traveled during the 5-min EPM session was not significantly different either (Fig. 4C; treatment effect: F_(5,57)_ = 0.8929, *p* = 0.4921; sex effect: F_(1,57)_ = 0.0405, *p* = 0.8412), indicating similar locomotor activity (Table 1).

**Figure 4.**
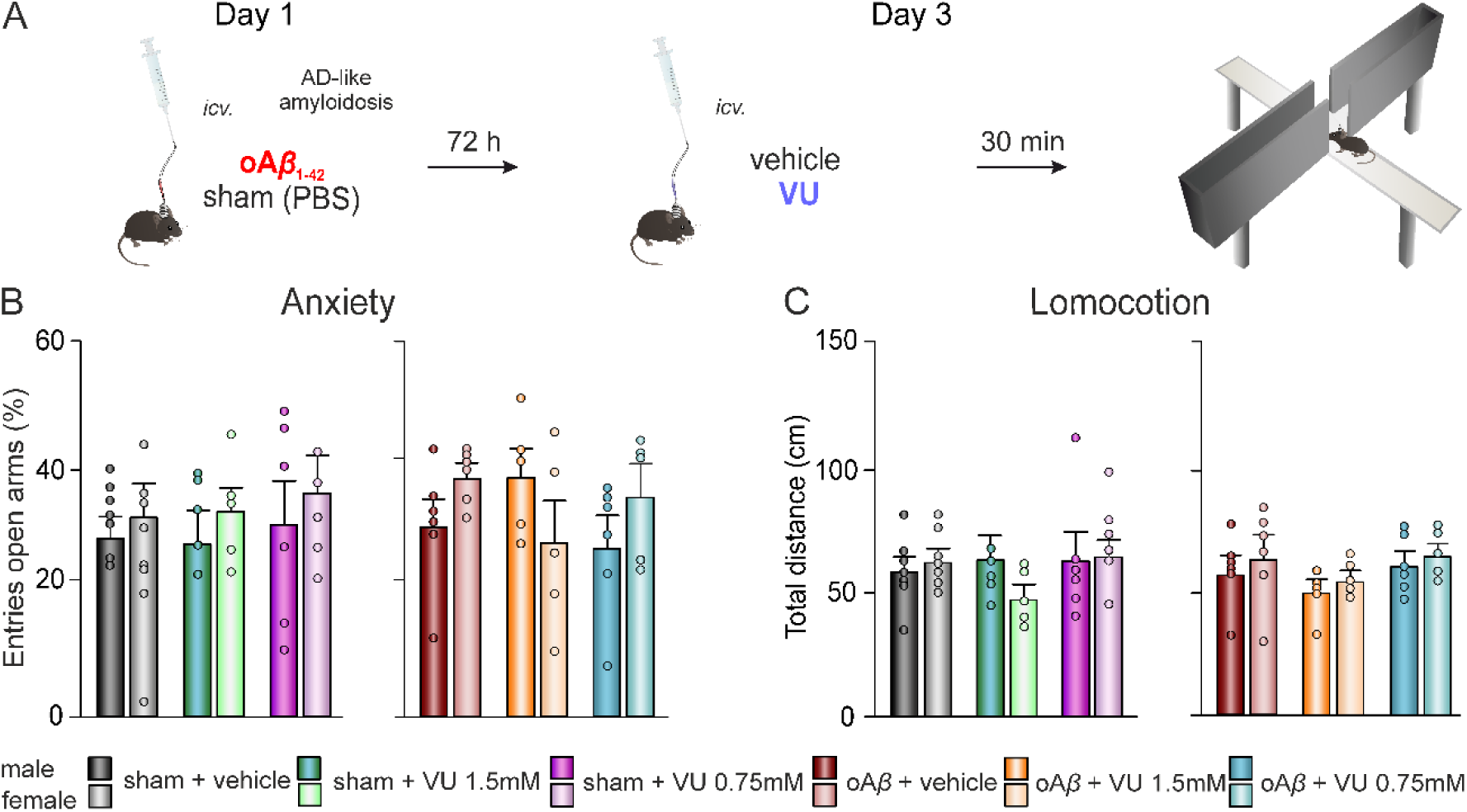
VU0810464 administration does not alter anxiety-like behavior and locomotion. **(A)** The EPM was used to evaluate the effect of VU0810464 (VU) treatment on the general health state of the animals, on day 3 post-oA*β*_1–42_ or sham (PBS) injection. **(B)** Percentage (%) of entries in open arms were measured to evaluate anxiety levels. **(C)** Total distance traveled was used as a measure of locomotor activity for (left) sham + vehicle, sham + VU 0.75mM and sham + VU 1.5mM treated mice and for (right) oA*β* + vehicle, oA*β* + VU 0.75mM and oA*β* + VU 1.5mM groups.

Thus, these data confirmed that overall health status and locomotor function were uniform among the different treated groups and, therefore, all learning and memory impairments observed in this work were primarily due to a specific hippocampal disruption caused by oA*β*_1–42_. Moreover, the improvement of those memory deficits can be attributed to the effect of VU0810464 on the hippocampus, and not to a potential drug influence on anxiety level or locomotion.

## 4. Discussion

GIRK channels have been recently revealed as key physiological determinant of neuronal excitability to support synaptic plasticity and related hippocampal-dependent cognitive functions in physiological conditions [19]. In addition, increasing evidence suggests a dysregulation in the signaling pathways mediated by, or converging on GIRK channels in AD [14, 18, 28, 30–32, 35]. Our results suggest that the precise tuning of neural excitability with low dosing of VU0810464, a novel GIRK activator, with greater potency and neuronal selectivity than previous compounds, might be a promising strategy to safely preserve hippocampal synaptic plasticity and upstreaming memory processes disrupted in early stages of AD.

### Early AD-like amyloidosis and sex dimorphism

Here, to further investigate the link between GIRK channels activity and AD, we used an acute animal model of AD generated by a single *icv*. injection of oA*β*_1–42_ that simulate the initial amyloidosis stage of the disease [40, 46], when hippocampal neuronal hyperexcitability initiates aberrant synaptic plasticity and patterns of neuronal circuit activity [10, 51], resulting in characteristic symptoms of MCI or early AD [12]. It has been previously shown that oA*β*_1–42_ *icv.* injection reaches both dorsal hippocampi generating an acute dorso-hippocampal amyloidosis which effects last for more than two weeks. One important advantage of this model is the induction of hippocampal cognitive deficits compatible with such AD stages in male and female mice [40]. Mice reach sexual maturity between 3 and 6 months of age and undergo their reproductive senescence transition between 9 and 12 months, marking the commonly characterized menopause, endocrine equivalent of the human period [52]. But menopause also include neurological symptoms involving disturbances in estrogen-regulated systems such as thermoregulation, sleep, and circadian rhythms, or may contribute to issues like depression and cognitive impairment [53]. The abrupt decrease in levels of estrogen and progesterone following menopause limits the neuroprotective actions that both sexual hormones exert on cerebral blood flow, neurotoxicity and hippocampal functionality [54]. In fact, increased A*β* burden has been shown in women who had experienced either natural or surgically induced menopause, with largest accumulation in those who underwent surgical menopause at a younger age [55]. In this context, our model allows the study of potential therapeutic targets in male and female mice before possible alterations related to the menopause period could contribute to the effects of oA*β*_1–42_.

### Pharmacological GIRK channels activation in health and disease

The strategy of regulating neural excitability to prevent subsequent upstreaming processes altered in AD (synaptic plasticity, oscillatory activity and behavior) has been gaining interest in the last decade based on evidence that subclinical epileptiform activity in patients with AD can also lead to accelerated cognitive decline. Treatment of clinical seizures in patients with AD with selective antiepileptic drugs, in low doses, is usually well tolerated and efficacious [56]. In fact, recent evidence of the cognitive improvement after treatment with low dose of levetiracetam in a visuospatial memory task, highly related to hippocampal function in AD patients, has been reported [57], although a larger sample would be required to test whether it might halt or reverse AD progression [58]. Supporting these hypotheses, studies in mouse models of AD consistently suggest that reducing network hyperexcitability have disease-modifying properties [59–62]. In this scenario, GIRK channels stay in a core position of hippocampal inhibitory neurotransmission, ruling the functions of different neurotransmission systems as downstream effectors of a variety of receptors [16], and therefore, becoming a potential target to normalize neural hyperactivity that characterize early AD [63, 64].

Regrettably, a major problem with classic selective GIRK channel activators is their poor pharmacokinetic profile. An example of that is ML297 mainly due to its urea-containing, which provides high hygroscopicity, poor solubility, low brain penetration, and modest selectivity for neuronal GIRK1/2 channel subtype [65]. To solve this issue, some authors have tried to dissolve it in saline +10% Tween-80 [50], or +10% Tween-20 [14], although in our hands, ML297 preparations resulted in a cloudy looking suspension. VU0810464 has been shown to present not only improved pharmacokinetic properties as a non-urea containing compound, but also selectivity, potency, and efficacy for GIRK channel activation, specifically in the brain, as well as higher blood brain barrier penetration [36, 44]. In addition, it has been recently proposed the use of HPβCD as an ideal excipient to improve its solubility. For intraperitoneal (*ip*.) administration, HPβCD has been used at 20% in sterile water, and led VU0810464 to stabilize as a microsuspension [44]. However, the route of administration in our experiments was *icv*., so HPβCD concentration was decreased to 10% in PBS in order to maintain stability of the solution, and preparation appeared crystalline. HPβCD has been previously used for *icv.* administration even at much higher concentration (40%) and volume (10µL) without evident effects on EEG activity or general behavior [66], and it is an approved excipient in both the United States and Europe, recommended as a solubilizer and stabilizer [67], so it seems an excellent adjuvant to facilitate VU0810464 *icv.* administration. Indeed, its *icv.* administration as vehicle in the present study did not induce any effect in control healthy or AD mice *in vivo*.

On the other hand, it has been previously reported that hippocampal GIRK activation with ML297 in physiological conditions impairs synaptic plasticity and oscillatory activity, leading to hippocampal dependent memory deficits [19, 25]. In these works, ML297 concentrations for *ex vivo* and *in vivo* experiments were 10 µM and 1.5 mM, respectively. Here, in order to evaluate whether the more selective VU0810464 presented the same beneficial effects on hippocampal-dependent memory, concentrations were half-reduced compared to ML297. Our data showed that *ex vivo* 10 µM remains significantly blocking LTP in control healthy animals, while a lower dose (5 µM) allowed to induce this synaptic plasticity process.

LTP, considered the cellular correlate of learning and memory [68, 69], has been systematically found to be altered *ex vivo* [28, 40, 70] and *in vivo* [14, 29] by early oA*β*_1-42_ forms. The more consistent explanation relies in the ability of oA*β*_1-42_ to promote neuronal hyperexcitability, directly disrupting neuronal receptors like NMDA, AMPA, and mGluRs, as well as astrocytic synaptic functions that altogether, impair hippocampal LTP and enhance LTD [10]. In fact, we previously found that a single *icv.* injection of oA*β*_1-42_ impairs LTP/LTD threshold when synaptic plasticity was measured from 1-17 days post-*icv.* [40]. This impairment in synaptic plasticity would explain the deficits observed in hippocampal-dependent spatial contextual recognition memory, tested by OLM [71], due to an altered threshold of LTP/LTD [47]. OLM seems to be mediated by hippocampal CA1 dopaminergic D1/D5 receptors [47] whose activation induce GIRK conductance to decrease [72]. If oA*β*_1-42_-induced hyperexcitation also impairs LTP/LTD threshold [28], GIRK channels would have serious difficulties to rule synaptic plasticity processes in the CA1 hippocampal region as previously found [19].

The correlation of *ex vivo* results at the behavioral level did show that higher concentration of VU0810464 (1.5 mM) blocked OLM in healthy, control animals, while 0.75 mM, the low one selected for *in vivo* experiments based on supplementary pilot study, had no significant effects on OLM performance. These results support the contention that using the appropriate dose of the drug, specifically a low dose, allows GIRK activity to effectively regulate synaptic plasticity processes, thereby sustaining hippocampal-dependent cognitive functions, which might have interesting clinical implications [37, 73]. However, high VU0810464 concentration disrupted OLM by hindering LTP. In this case, hyperpolarization due to enhanced GIRK conductance might explain why the HFS protocol fails to sufficiently activate NMDA receptors, leading to a slight rise in intracellular Ca^2+^ levels and subsequently causing a decrease in synaptic efficacy [74]. However, as previously mentioned, its high selectivity and fast clearance [44] may account for the induction of a lower depression compared to that caused by ML297 [19]. Nonetheless, in a pathological context like AD where the E/I balance is disrupted, the corrective effects of stimulating GIRK channels at the synaptic level, with both concentration of VU0810464, could prompt the restoration of the excitability levels impaired by oA*β*, and subsequent synaptic plasticity and memory processes.

We also evaluated the possible GIRK-dependent decrease in anxiety-related behavior previously described using the EPM test when ML297 is *ip.* administered [50]. This is an important issue to consider when behavioral tests, such as OLM, are performed, as anxiety might affect results interpretation. However, evidence has shown that ML297 *icv*. injected was not associated with abnormal anxiety levels [19]. Likewise, VU0810464 does not impact EPM performance when *ip*. administered [44] and here, *icv*. injections had no effects on anxiety levels neither. One plausible explanation for the differential behavioral outcomes of GIRK activation in the EPM test could result from the faster clearance of VU0810464 compared to ML297, a determinant fact for its potential broad utility *in vivo*, although it could also be related to the selectivity for channel subunit(s) involved in this behavior [44].

In summary, the results obtained in this work underpin the relevance of low dosing of VU0810464 -an easy-soluble and fast-cleared, highly selective, potent and effective neuronal GIRK1/2 channel activator-as a potential pharmacological manipulation to counteract early AD-related neuronal excitability disruptions and improve hippocampal synaptic plasticity and dependent memory before potential sex differences associated with age become evident. But which is more crucial, targeting neuronal GIRK channels without the significant secondary effects of enhancing their activity [19, 25] in physiological conditions.

## 5. Conclusion

Recent evidence has shown that monotherapy regimen with low-dose of anti-seizure drugs can improve outcomes in AD patients with epileptic activity [73] and may have disease-modifying effects involving A*β* and tau [56]. But it also suggests that treatments should be used in the prodromic stages of AD, in combination with identification of new biomarkers to promote multi-target personalized therapy for individuals in the earliest stages of neurodegenerative diseases [75]. In this framework, our results are particularly relevant for investigators that are seeking effective therapies to chronically treat epileptic activity in the form of early hyperexcitability, associated with AD [37]. VU0810464 might represent a potential new drug highly selective on neuronal GIRK1/2 channels with favorable pharmacokinetic and pharmacodynamic properties [44], to normalize cognitive and functional outcomes by preventing the presence of excess neural activity, a pathological hallmark of early AD. Furthermore, a high concentration of VU0810464 significantly blocked synaptic plasticity and impaired hippocampal-dependent memory in healthy animals, while the low concentration did not have these undesirable effects. This suggests that the latter low dose could be a potential treatment not only for early AD but also as a preventive therapy for the preclinical stages of this dementia, although further experiments are needed to validate this hypothesis.

## Declarations

### Data availability statement

All data supporting the results in the paper are in the paper itself and the supporting information. Raw data and recordings are available in the laboratory from the corresponding authors and will be provided upon reasonable request.

### Ethics approval

All experimental procedures were reviewed and approved by the Ethical Committee for Use of Laboratory Animals of the University of Castilla-La Mancha (PR-2021-12-21 and PR-2023-25) and conducted according to the European Union guidelines (2010/63/EU) and the Spanish regulations for the use of laboratory animals in chronic experiments (RD 53/2013 on the care of experimental animals: BOE 08/02/2013).

### CRediT authorship contribution statement

LJD and JDNL were responsible for the initial conceptualization; JMF, RJH, AC, and SD performed methodology, data curation and formal analysis; JMF and RJH were responsible for the visualization; JDNL was responsible for writing the original draft; JDNL, AC, SD and LJD did the writing – review and editing. LJD and JDNL were responsible for funding acquisition, project administration and supervision. All authors read and approved the final manuscript.

### Declaration of Competing Interest

The authors declare that they have no known competing financial interests or personal relationships that could have appeared to influence the work reported in this paper.

### Funding sources

This work was supported by MCIN/AEI/10.13039/501100011033 (grant number PID2020-115823-GBI00), JCCM and ERDF A way of making Europe (grant number SBPLY/21/180501/000150) and 2022-GRIN-34354 grant by UCLM/ERDF intramural funds to JDNL and LJD; JMF held a predoctoral fellowship granted by JCCM and ERDF-A way of making Europe Program (2023-PRED-21780). AC held a *Margarita Salas* Postdoctoral Research Fellow (2021-MS-20549) funded by European Union NextGenerationEU/PRTR.

## Acknowledgements

We are deeply grateful to Prof. Javer Yajeya, who, although no longer with us, continues to inspire this research line.

## Supplementary file 1

### Supplementary materials and methods

#### Drug dosage

To determine the potential side effects of *in vivo* VU0810464 administration and therefore choose the most appropriate dosage to further study it as an acute amyloidosis treatment, an extensive set of behavioral tests was carried out.

Thus, a cohort of 29 C57BL/6 adult mice (14 females and 15 males; 3 -6 months old; 20 - 35 g) were used, and randomly divided into three groups: VU0810464 0.25mM, VU0810464 0.75mM and Veh (PBS with 10% HPβCD).

Animals receive a daily *icv.* injection (1µl) of the corresponding treatment for 3 consecutive days 30 min prior to the OLM, EPM and rotarod tests, then a 10-day period of recovery was allowed to prevent dilatation of the lateral ventricle. Afterwards, daily *icv.* injections were repeated for another 3 days prior to the open field (OF) habituation, stereotyped and locomotion and tail suspension tests, followed by another 10-day recovery period. Finally, another OF was performed to confirmed that the drug was cleared before starting the Barnes maze (Suppl. Figure S1A).

Both the OLM (Suppl. Figure S1B) and the EPM (Suppl. Figure S1C) were performed as previously described in the main text.

#### Rotarod performance test

To assess motor coordination and learning, mice were trained to stay for 1 min at constant low-speed (6 rpm) rotation on the rotarod apparatus (LE 8500, Panlab, Spain), which consists of a 30 mm diameter black striated rod positioned 20 cm above the floor. Then, 5 consecutive trials with an acceleration from 4 to 40 rpm over a 2 min period were carried out, measuring mice’s time to fall off the rod (Suppl. Figure S1D).

#### Open field habituation task

To evaluate a non-associative hippocampal-dependent learning process, an OF habituation task was performed, in which animals were exposed to a novel arena on two consecutive days and the change in exploration time after re-exposure was measured. Thus, mice were place in a square acrylic box (23.5 × 17.5 × 4 cm plexiglas base arena; 26.5 × 21 × 10 cm top) and allowed to freely explore for 15 min during the training day (OF1) and 24h later during the retrieval day (OF2) (Suppl. Figure S1E and 1H).

#### Laboratory animal behavior observation registration and analysis system (LABORAS®) for stereotyped and locomotion behavioral testing

To evaluate stereotyped behaviors and locomotion, a LABORAS® apparatus (Metris, The Netherlands) was used, which captures mechanical vibrations generated by the movements of the animals and convert them into electrical signals through a sensing platform positioned beneath the cage. Mice were placed in the LABORAS® cage for a single 15-min trial, and locomotion, climbing, rearing and grooming were recorded as measured of locomotor activity, exploration and stress-related behaviors (Suppl. Figure S1F).

#### Tail suspension test

To assess depression-like behaviors, the tail suspension test (BIO-TST5, Bioseb, US) was performed, which consisted in three PVC chambers (50 × 15 × 30 cm each) in which animals were hung by their tails approximately 10 cm above the ground for 6 min. Immobility time, energy and power were recorded using strain sensors and digitized for further analysis using the BIO-TST5 software (Bioseb, USA) (Suppl. Figure S1G).

#### Barnes maze

To evaluate hippocampal-dependent working, short-term and long-term spatial memory, the Barnes maze (LE851BSW, Panlab, Spain) was performed with a protocol previously used [1]. Briefly, in each trial the mice were placed inside a starting cylinder (8 × 12.5 cm) in the center of a platform disk (92 cm in diameter) with 20 escape holes (5 cm each) around the periphery, elevated 1 m above the floor and with spatial cues around the room’s walls. After 10 sec, the cylinder was removed, a mildly aversive white noise was started, and animals were allowed to explore until they find the escape box (17.5 × 7.5 × 8 cm) attached to one of the holes or the time limit was reached. In both cases, mice remained in the box for 1 min before being returned to their cages. Between each trial, the maze was cleaned with 70% ethanol to dissipate odor cues.

The protocol consisted in a habituation day, 4 training days and 2 memory tests days. On training days 1 and 2, animals did not receive any treatment, to make sure that the learning process was achieve equally in all groups. Afterwards, animals were *icv.* injected 30-min prior training days 3 and 4, as well as the two memory tests.

Habituation consisted in a single habituation trial of 90 s performed 24 h before training. Then, training was carried out over 4 consecutive days, with 3 trials per day of 3 min each and a 15 min interval between trials. During training, the escape hole changed daily, and latency, distance, and probability of success were measured. Finally, each memory test day a single 90 s trial was performed, with no escape hole available, and latency, distance and probability of reaching the target of the latest training day were measured (Suppl. Figure S1I).

All sessions were recorded and analyzed with Barnes-Smart video tracking software (Panlab, Spain). The cumulative incidence of latency to find the escape hole was analyzed using a Cox proportional hazard model, with group as a categorical covariate and day as continuous covariate.

### Supplementary results

Previously, our group had proved that both gain and loss of GIRK-channel function had deleterious effects on synaptic plasticity and memory in healthy, naïve mice, using the blocker Tertiapin-Q and the opener ML297 [2]. Nevertheless, VU0810464 seems to be a more specific and potent GIRK opener that ML297, therefore potentially been a better candidate as a treatment for acute amyloidosis while avoiding adverse effects. Thus, before testing the effect of this drug in a non-transgenic model of AD, a pharmacological characterization was carried out to determine a dose that would not cause adverse effects, comparing the *icv.* administration of 0.25mM and 0.75mM of VU.

#### Effect of VU on motor activity

Mobility, motor coordination and motor learning were studied using a range of tests.

Firstly, mobility was examined with an object location memory test (Suppl. Figure S1B). During the habituation session, before any treatment was administered, distance travel was measured to ensure that all naïve animals had a similar behavior. Indeed, as shown in Suppl. Figure S1B, all animals traveled on average the same distance (F_(2,15)_ = 2.103, *p* = 0.1566). However, when analyzing the distance traveled after a single *icv*. VU injection, results showed that mice moved less specifically in the OLM1 when treated with VU at a concentration of 0.25 mM (F_(2,15)_ = 5.816, *p* = 0.0135).

Mobility was also measured using the elevated plus maze (Suppl. Figure S1C), and in this test it was not altered in any treatment group, since all showed similar distance traveled (F_(2,24)_ = 0.3427, *p* = 0.7133).

However, when comparing the distance traveled in a OF habituation task (Suppl. Figure S1E), a significant treatment effect was found (F_(2,25)_ = 3.662, *p* = 0.04), showing lower mobility in the VU 0.25mM group compared to both the control and the VU 0.75mM groups during the OF1.

Since mobility seems to be altered, motor coordination and learning were then evaluated using the rotarod performance test (Suppl. Figure S1D). In line with the previous results, motor coordination was impaired after VU 0.25mM treatment since they showed a lower latency to fall of the rod than vehicle animals (t_18_ = 2.033, *p* = 0.05, *p* = 0.7133). However, all groups improved their performance along trials, showing proper motor learning (Time effect: F_(3.829,99.55)_ = 2.702, *p =* 0.0369, Geisser-Greenhouse’s correction; Treatment effect: F_(2,26)_ = 1.17, *p* = 0.2).

#### Effect of VU on stereotyped behavior

To further assess the overall health of the animals after treatment with different concentrations of VU, a wide range of physiological behaviors were analyzed in a LABORAS apparatus (Suppl. Figure S1F).

One-way ANOVAs revealed significant differences between VU 0.25mM mice and the other two groups in locomotion (F_(2,26)_ = 3.424, *p* = 0.0479), immobility (F_(2,25)_ = 4.351, *p* = 0.0239), rearing (F_(2,26)_ = 9.203, *p* = 0.001) and grooming (F_(2,26)_ = 5.275, *p* = 0.0119). Locomotion and rearing, a behavior related to the exploratory nature of mice, were reduced, while immobility and grooming were enhanced. No statistical difference was found in climbing (F_(2,25)_ = 1.638, *p* = 0.2146).

#### Effect of VU on anxiety-and depression-like behaviors

To assess anxiety-like behaviors, the elevated plus maze was used the second day of *icv.* injection (Suppl. Figure S1C), and data showed no altered anxiety levels in any treatment group, since all showed similar entries into the open arms (F_(2,24)_ = 0.4643, *p* = 0.6341).

When studying depression-like behavior using the tail suspension test (Suppl. Figure S1G), no differences among any of the groups were found, neither in the immobility time (F_(2,25)_ = 1.775, *p* = 0.19) nor in the energy or the power of the movements (F_(2,23)_ = 0.9657, *p* = 0.3956; F_(2,23)_ = 2.185, *p* = 0.1352, respectively).

#### Effect of VU on spatial memory

The Object location memory test was used to study spatial memory (Suppl. Figure S1F). Although animals showed an alteration in mobility, this did not affect spatial memory, as all groups showed equally robust discrimination index in both short (OLM1) and long term (OLM2) memory (Time effect: F_(1.66,24.9)_ = 23.29, *p* < 0.0001, Geisser-Greenhouse’s correction; Treatment effect: F_(2,15)_ = 0.387, *p* = 0.6857).

Later on, a far more complex memory test such as the Barnes maze was carried out. Before that, animals underwent another OF (Suppl. Figure S1H), ten days after the last *icv*. injection of VU, to ensure that the treatment was cleared out and all animals were back to baseline levels regarding their mobility. Indeed, all groups showed equal intrasession learning (Time effect: F_(6.477,168.4)_ = 34.84, *p* < 0.0001, Geisser-Greenhouse’s correction) and no difference among them in distance traveled (Treatment effect: F_(2,26)_ = 0.8123, *p* = 0.4548).

Thus, the Barnes maze was applied in “naïve” animals. First, the distance traveled during the habituation session (before treatment) was measured and no differences between groups were found (Suppl. Figure S1I; F_(2,26)_ = 2.416, *p* = 0.109).

Latency to find the escape hole was assess as a measure of memory (Suppl. Figure S1I), and two-way ANOVA showed that all groups improved their latency along days (Time effect: F_(3.058,77.07)_ = 77.07, *p* < 0.0001, Geisser-Greenhouse’s correction; Treatment effect: F_(2,26)_ = 0.9629, *p* = 0.395). Although no treatment effect was found, there was a significant difference (t_18_ = 2.441, *p* = 0.0252) between VU 0.25mM and vehicles on day 3 of training (i.e., the first day of *icv*. injection).

Accordingly, the distance covered gradually decreased over the course of the training sessions and tests as the animals learned to perform the task (Suppl. Figure S1I; Time effect: F_(3.621,91.98)_ = 44.87, *p* < 0.0001, Geisser-Greenhouse’s correction; Treatment effect: F_(2,26)_ = 0.6275, *p* = 0.5418).

Furthermore, a Cox proportional hazard model was applied to estimate the extent of the effect that the treatment had on the probability of successfully completing the task (i.e. finding the escape hole) [3]. Data showed that both treatment and time had a significant effect on the success rate (Suppl. Figure S1I), with the probability of reaching the escape hole being decreased by 33.2% after VU 0.25mM administration (Hazard ratio = 0.668, 95% CI = 0.461-0.968, *p* = 0.033) but not after VU 0.75mM (*p* = 0.466). Moreover, across all groups this rate improved by 50% with each day (Hazard ratio = 1.50, 95% CI = 1.395-1.613, *p* < 0.001).

#### Effect of VU on non-associative memory

To assess a hippocampal-dependent non-associative type of memory, the open field habituation test was performed.

The learning process seemed to be altered in the VU 0.25mM group, as appraised by the analysis of the intrasession (Suppl. Figure S1E; Time effect: F_(6.48,162)_ = 71.25, *p* < 0.0001, Geisser-Greenhouse’s correction; Treatment effect: F_(2,25)_ = 4.206, *p* = 0.0266). Nevertheless, memory was undisturbed, since all groups traveled less distance in the OF2 compared to their corresponding OF1 (Suppl. Figure S1E; Time effect: F_(1,25)_ = 41.62, *p* < 0.0001).

Overall, all the data presented above indicate that the 0.25mM dose of VU is a poor candidate to be used as a treatment for acute amyloidosis since, although it does not appear to affect memory in healthy animals, it does cause noticeable side effects on their mobility, motor coordination and stereotypes behaviors. However, the 0.75mM dose did not cause any of those adverse effects in healthy animals, so we set out to further study it to determine whether it could reverse the effects of A*β* in our non-transgenic mice model of AD.

---

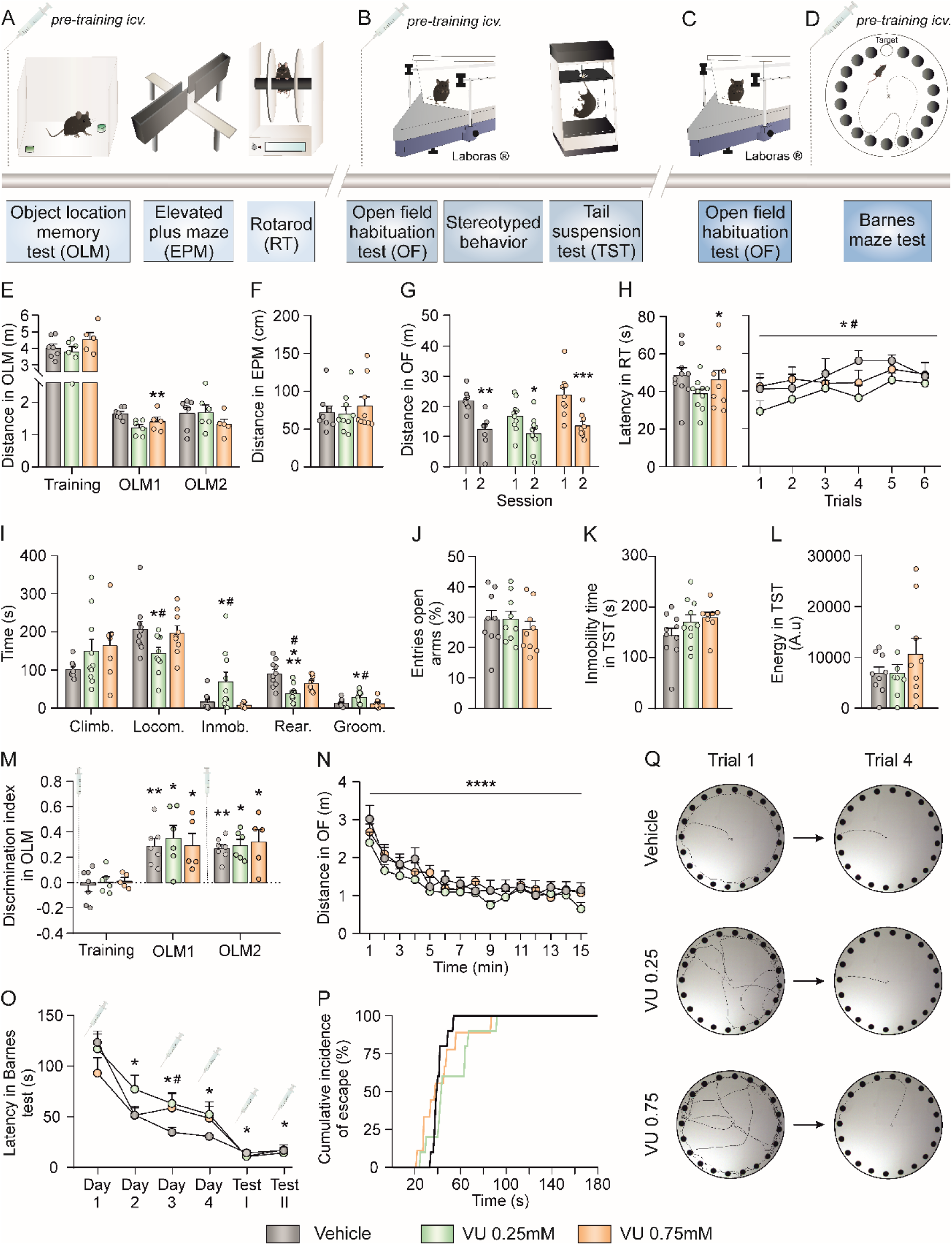

## Notes

### Competing Interest Statement

The authors have declared no competing interest.

